# Optoacoustic brains stimulation at submillimeter spatial precision

**DOI:** 10.1101/459933

**Authors:** Ying Jiang, Hyeon Jeong Lee, Lu Lan, Hua-an Tseng, Chen Yang, Heng-Ye Man, Xue Han, Ji-Xin Cheng

## Abstract

Low-intensity ultrasound is an emerging modality for neuromodulation. Yet, piezo-based transducers offer poor spatial confinement of excitation volume, often bigger than a few millimeters in diameter. In addition, the bulky size limits their implementation in a wearable setting and prevents integration with other experimental modalities. Here, we report spatially confined optoacoustic neural stimulation through a novel miniaturized Fiber-Optoacoustic Converter (FOC). The FOC has a diameter of 600 μm and generates omnidirectional ultrasound wave locally at the fiber tip through the optoacoustic effect. We show that the optoacoustic wave can directly activate individual cultured neurons and generate intracellular Ca^2+^ transients. The FOC activates neurons within a radius of 500 μm around the fiber tip, delivering superior spatial resolution over conventional piezo-based low-frequency transducers. Combining FOC with electrophysiology, direct and spatially confined neural stimulation of mouse brain is achieved *in vivo.*

## Main

Ultrasound is an emerging neuromodulation technique that offers the potential of non-invasively modulating brain activities ^1,2^ Early reports of neuromodulation using high-intensity ultrasound date back to the 1920s, likely through a tissue heating mechanism^3,4^. In the past decade, neural stimulation using low intensity, low frequency focused ultrasound has been demonstrated to directly evoke action potentials and modulate motor response in rodents^2,5^, rabbits^6^, non-human primates^7^ and sensory/motor responses in humans^8-11^ through a non-thermal mechanism. Two recent papers argued that these responses could be a consequence of indirect auditory stimulation through the cochlear pathway^12,13^. Yet, many others have reported direct activation of neurons in brain slices^14^ and isolated retina^15,16^, where no auditory circuitry is involved. A major challenge facing ultrasound neural modulation, which contributes to the mentioned controversies, is that delivery of transcranial ultrasound would inevitably go through the skull, and eventually reach the cochlear through bone transduction. Moreover, the presence of the skull will reflect acoustic wave and compromise ultrasound focus, resulting in poor spatial resolution. l

An alternative way to generate ultrasound wave is through optoacoustic effect. In an optoacoustic process, pulsed light is illuminated on an absorber, causing transient heating and thermal expansion, and generating broadband acoustic waves at ultrasonic frequencies^17^. Recently, the optoacoustic effect has received increasing attention in the fields of imaging and translational medicine^18,19^. Using endogenous as well as exogenous absorbers, optoacoustic tomography and microscopy have found broad biomedical applications^20,21^. Beyond imaging, recent advances in developing optoacoustic materials have enabled highly efficient optoacoustic conversion ^22^. Pulsed light excitation of these optoacoustic materials generates ultrasound waves at high amplitude, which allowed for all-optical ultrasound imaging ^23,24^, tissue cavitation ^25,26^, and precision surgical guidance of lumpectomy ^27^.

Here, we report the first use of optoacoustic wave for direct and spatially confined neural stimulation both in culture and in the living brain. The stimulation is based on a novel fiber optoacoustic converter (FOC) that generates omnidirectional ultrasound pulses emitting from a coated fiber tip. The miniaturized size of the FOC together with fast ultrasound attenuation provides superior spatial confinement of the generated ultrasound. By time-resolved calcium imaging, we demonstrate that the FOC can reliably produce neural activation within a 500 μm radius from the FOC tip in cultured neurons, and the stimulation effect is specific to neurons. By combining FOC with electrophysiology, we achieved direct optoacoustic activation of mouse somatosensory cortex in the living brain, providing evidence that the observed activation is a consequence of direct neural stimulation without the involvement of the cochlear pathway.

## Results

### Fabrication and characterization of FOC

The FOC is composed of a compact 1030-nm, 3-nanosecond laser, a 200-μm diameter, 0.22 NA multimodal fiber, and a ball-shaped coated tip with a diameter of ∼600 μm (**Fig. 1a**). Through the optoacoustic process, the pulsed laser energy is converted into acoustic waves generated at the FOC tip. The acoustic waves then excite neurons in the proximity to the tip. The FOC tip was coated with 2-layer nano-composite (**Fig. 1b**). The first layer is a diffusion layer composed of a mixture of ZnO nanoparticles in epoxy (10% w/w). The ZnO nanoparticles have a 100-nm diameter, which is smaller than the wavelength of the incident light and enables Raleigh scattering of the light. Consequently, the incident light is randomly scattered in all directions, which produces a relatively uniform angular distribution of the laser pulse. The second layer is an absorption layer composed of a mixture of graphite powders in epoxy (30% w/w). With its high optical absorption and thermal conduction efficiency, the graphite completely absorbs the diffused laser and converts it into heat. The heat is then transferred to surrounding epoxy, creating expansion and compression of the epoxy, and generating optoacoustic waves that propagate in an omnidirectional manner. To characterize the generated optoacoustic wave from FOC, we applied the nanosecond laser at a pulse energy of 14.5 μJ, and measured the optoacoustic wave by a needle hydrophone under water. A representative acoustic wave generated by a single laser pulse is shown in **Fig. 1C**. The radiofrequency spectrum shows that the generated acoustic wave is in the ultrasound frequency ranging from 0.5 to 5 MHz, with multiple peaks between 1 and 5 MHz (**Fig. 1d**). The maximum acoustic pressure is measured to be 0.1 Mega Pa. To examine the angular distribution of the optoacoustic wave, we measured the pressures at various angles. The distribution map shows that the intensity is the strongest in the forward direction, while the back-propagating ultrasound is about 50% of the forward intensity (**Fig. 1e**). The omnidirectional acoustic wave propagation is achieved by the diffusion layer and the ball-shaped geometry of the FOC, which allows the acoustic intensity to attenuate quickly when propagating in the medium.

**Figure 1.**
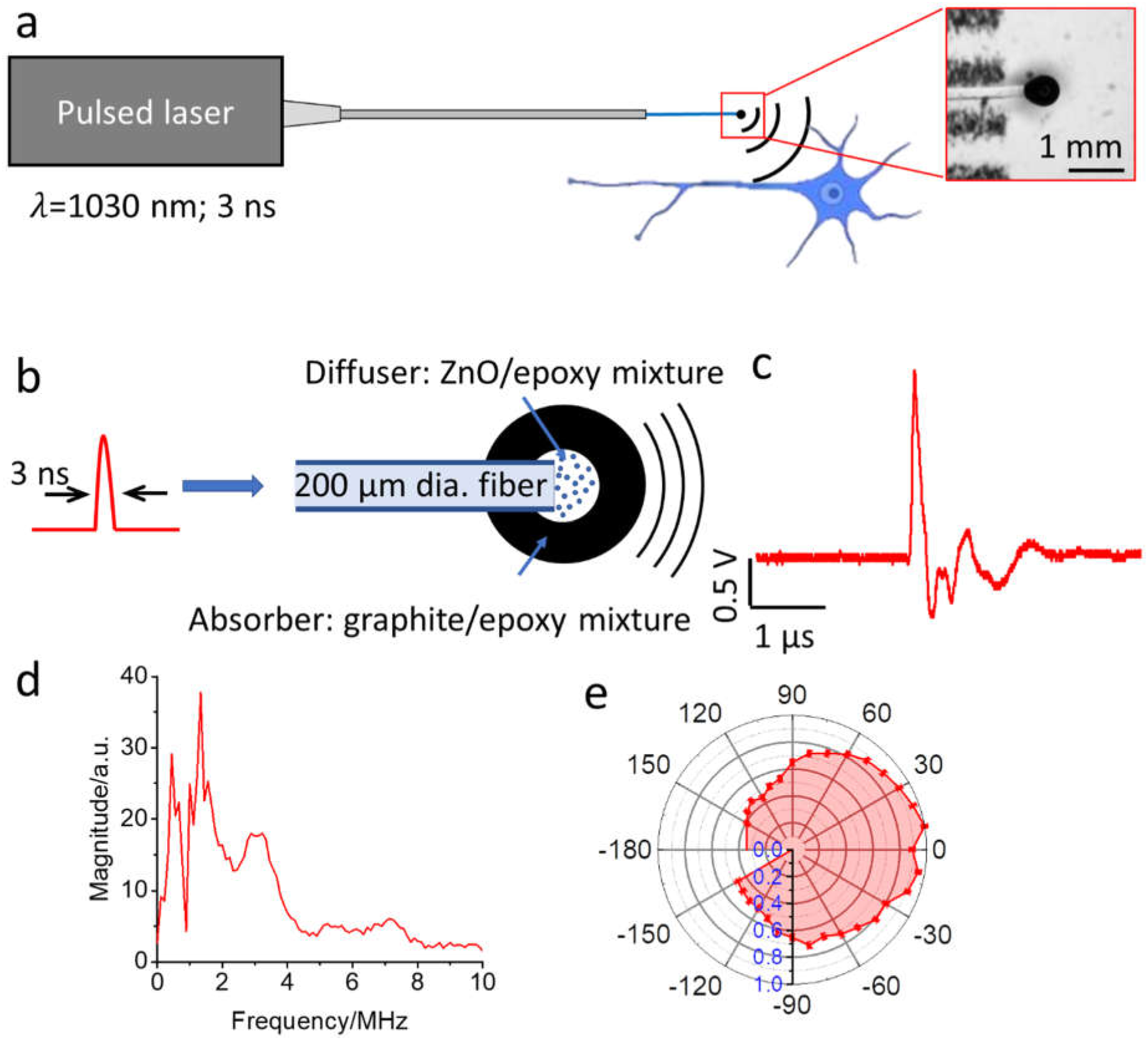
Design of FOC and characterization of the FOC-generated acoustic wave. **a.** The concept of optoacoustic neuromodulation through a FOC. Insert is an enlarged FOC tip under a stereoscope. **b**, Schematic of optoacoustic wave generation. **c**, Representative optoacoustic wave recorded with a hydrophone. **d,e**, radiofrequency spectrum and angler distribution of FOC generated acoustic wave. Error bar: ± SD

### FOC induces calcium transients in cultured neurons with high spatial precision

To investigate whether the FOC can directly modulate neuronal activity, we examined the FOC invoked response in cultured neurons. We treated rat cortical neurons (Days in *vitro* 18 to 22) with a calcium indicator, Oregon Green™ 488 BAPTA-1, AM (OGD-1), (**Fig. 2a**) and performed calcium imaging using an inverted wide-field fluorescence microscope (Fig. S1). The FOC was placed approximately 100 μm above the focus plane, in the center of the field of view. Optoacoustic wave is produced with a 200 ms long, 3.6 kHz laser pulse, which corresponds to approximately 720 acoustic pulses. Calcium transients were observed for all neurons in the field of view (max ΔF/F = 9.5 ± 2.9%, 36 cells from 3 cultures) (**Fig. 2b**). Addition of 15 μM intracellular calcium chelator BAPTA-AM significantly reduced the calcium signal (max ΔF/F = 2.6 ± 0.5%, n=12) (Fig. S2). The response latency was found to be less than 50 ms, since the responses were observed at the first frame post stimulation onset across experiments with a camera acquisition rate at 20 Hz. To identify the threshold for FOC induced neural activation, we varied the stimulation duration to 100, 50 and 20 ms. The FOC successfully produced neural activation with 100 and 50 ms stimulation, but not with 20 ms stimulation (**Fig. 2c**). We next asked whether the FOC can produce neural activation reliably and repeatedly. Eight burst of laser pulses, each with 200 ms burst duration and 2-second inter-burst interval, were delivered to the FOC. Stable calcium transients in response to each laser pulse train were observed (max ΔF/F = 12 ± 0.6%, 8 pulses), (**Fig. 2d**). No obvious morphological changes were detected in neurons stimulated multiple times over a 2-hour duration (**Fig. 2e**).

**Figure 2.**
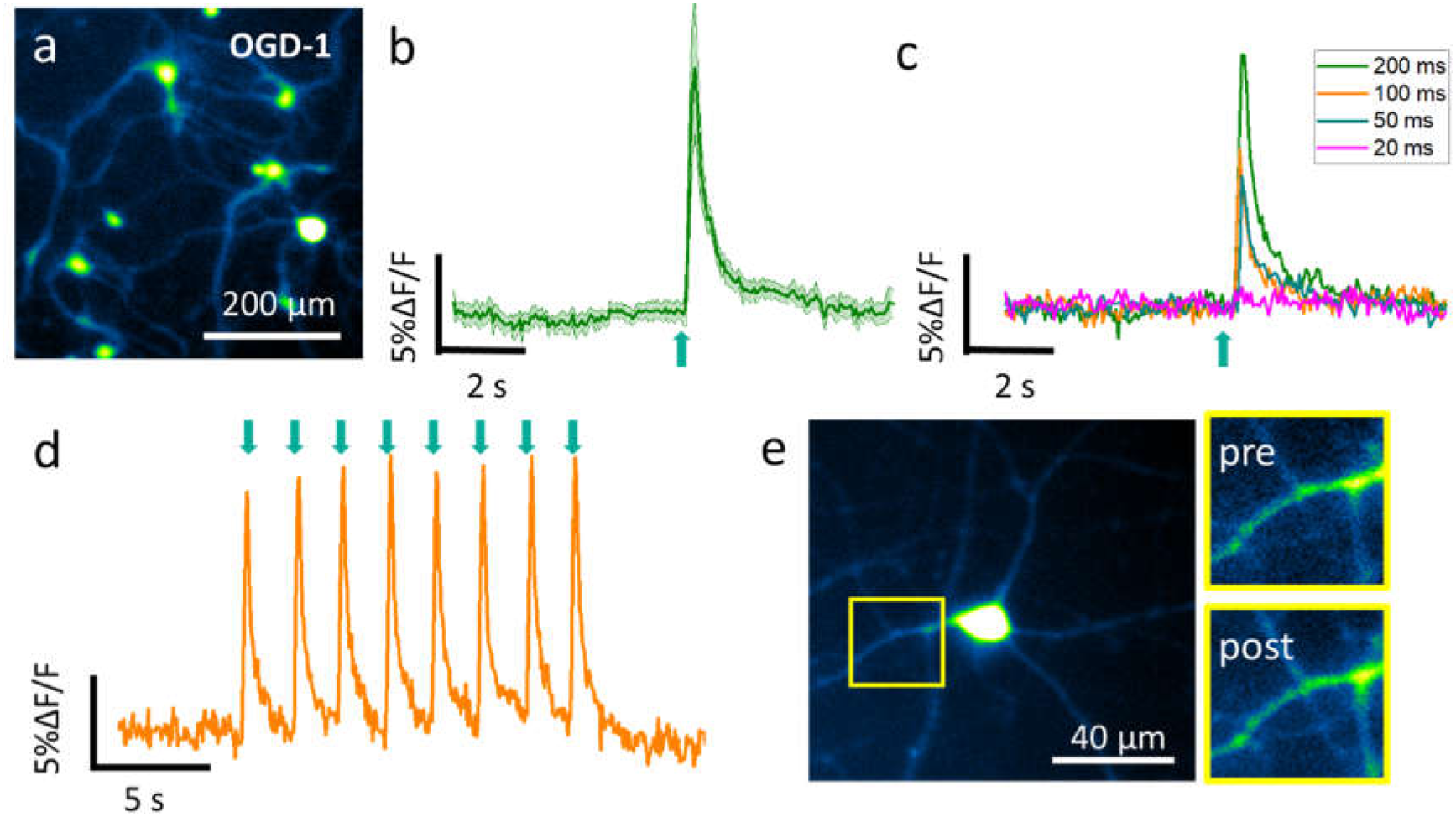
Optoacoustic wave induces calcium transients in cultured primary neurons. **a**, Neurons loaded with OGD-1. **b**, The average trace of neuronal calcium trace (n=12) in response to 200 ms FOC stimulation. Shaded area: ± SD. **c**, Representative traces of neuronal response to 200-ms, 100-ms, 50-ms, and 20-ms FOC stimulation. **d.** Calcium trace of a neuron undergone repeated FOC stimulations. **e.** Representative image of a neuron before and after repeated FOC stimulations. Green arrows: stimulation onset.

To ask whether the FOC-induced calcium transients is specific to neurons, we loaded OGD-1 to a prostate cancer cell line (PC3) and observed no calcium signal induced by 200-ms FOC stimulation (max ΔF/F = 0.7 ± 2.7%, n=52) (Fig. S3). Next, we obtained a rat glial culture loaded with OGD-1 and delivered 200-ms FOC stimulation to morphologically identified astrocytes, and observed responses (max ΔF/F = 1.2 ± 0.6%, n=82) significantly smaller than the calcium transients produced by FOC stimulated neurons (Fig. S4). The glial culture was immuno-stained after experiment and was confirmed as GFAP positive. These data suggest that the FOC reliably and selectively activates neuronal cells in vitro.

A key advantage of FOC over traditional ultrasound transducers is that the FOC emits pulsed ultrasound waves locally at the coated fiber tip, which allows localized stimulation. With a FOC tip of 600 μm in diameter, the acoustic intensity is attenuated by 56% at 1 mm away from the tip under water, measured by a transducer array followed with wavefront reconstruction (**Fig. 3 a,b**). Since the brain tissue has much higher ultrasound attenuation coefficient (0.6 dB/cm MHz) than water (0.0022 dB/cm MHz)^28^, we expect the FOC to produce even more localized acoustic wave in the brain. To demonstrate that FOC induced neural activation is spatially confined, we placed the FOC at the edge of the imaging field of view and delivered a laser pulse train of 200 ms duration (**Fig. 3c**). It was observed that neurons within 500 μm distance from the FOC showed reliable calcium response, and that the amplitude of the response is highly dependent on the relative distance from the FOC (**Fig. 3d**). When we sorted the neurons by their distance from the FOC, we found neurons that are closest to the FOC showed the largest response, while neurons that are 500 μm to 1.0 millimeter away showed negligible response (**Fig. 3e**). These data demonstrated that the effect of FOC is highly localized within a 500 μm radius, which provides one order of magnitude better spatial resolution comparing to conventional transducers with several-millimeter focus area.

**Figure 3.**
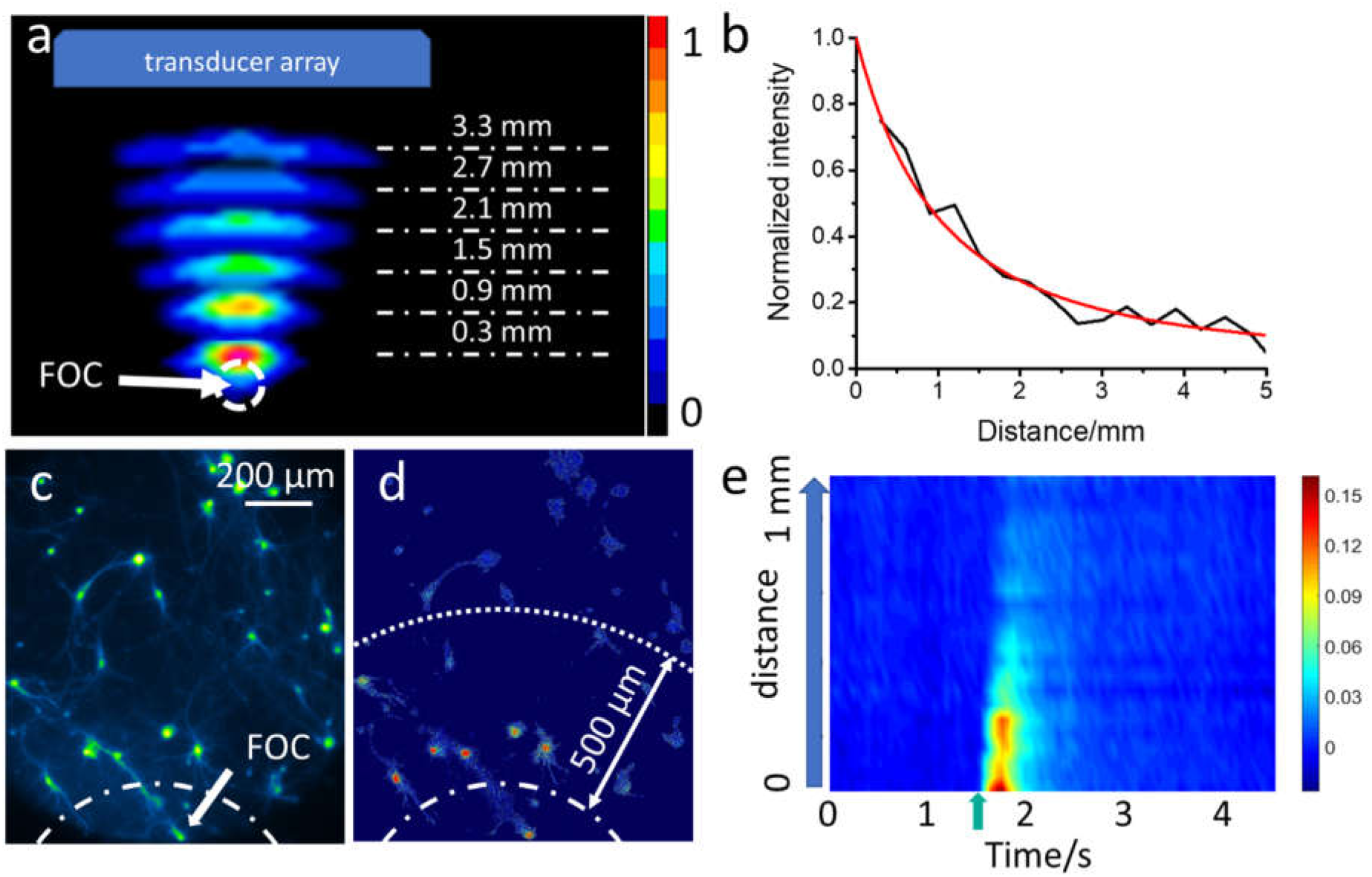
Spatially confined acoustic wave and neural stimulation induced by FOC. **a**, FOC wavefront reconstruction by the transducer array. Note only part of the wavefront is reconstructed due to the limited receptive angle of the transducer array. **b**, Acoustic intensity attenuates significantly as the distance to the FOC increases. **c,d**, Spatial distribution of maximum neuronal calcium response induced by 200 ms FOC stimulation. Dashed line: placement of FOC. **e**, Sorted neuronal calcium traces in relation to the distance from the FOC. Green arrow: stimulation onset.

To eliminate the possibility that the activation is due to laser illumination, we measured the leaked light energy from the FOC tip with a photodiode and found only 0.11% of the laser leaked out of the FOC. Additionally, we used an uncoated optical fiber and delivered the same laser pulses at 3.6 KHz repetition rate and 200 ms duration directly to the neuron. No calcium transients were observed (**Fig. 4a**). To examine the possibility of photothermal neural activation, we measured the heat profile of the FOC tip using a miniaturized ultrafast thermal probe. The temperature increase on the FOC surface was found to be 1.6, 0.9, 0.5 °C for 200, 100, 50 ms laser stimulations, respectively (**Fig. 4b**). Such temperature increase is well below the previously reported threshold for thermal induced neural activation (ΔT > 5 °C)^29^. Therefore, the effect of FOC is most likely contributed by the generated optoacoustic wave.

**Figure 4.**
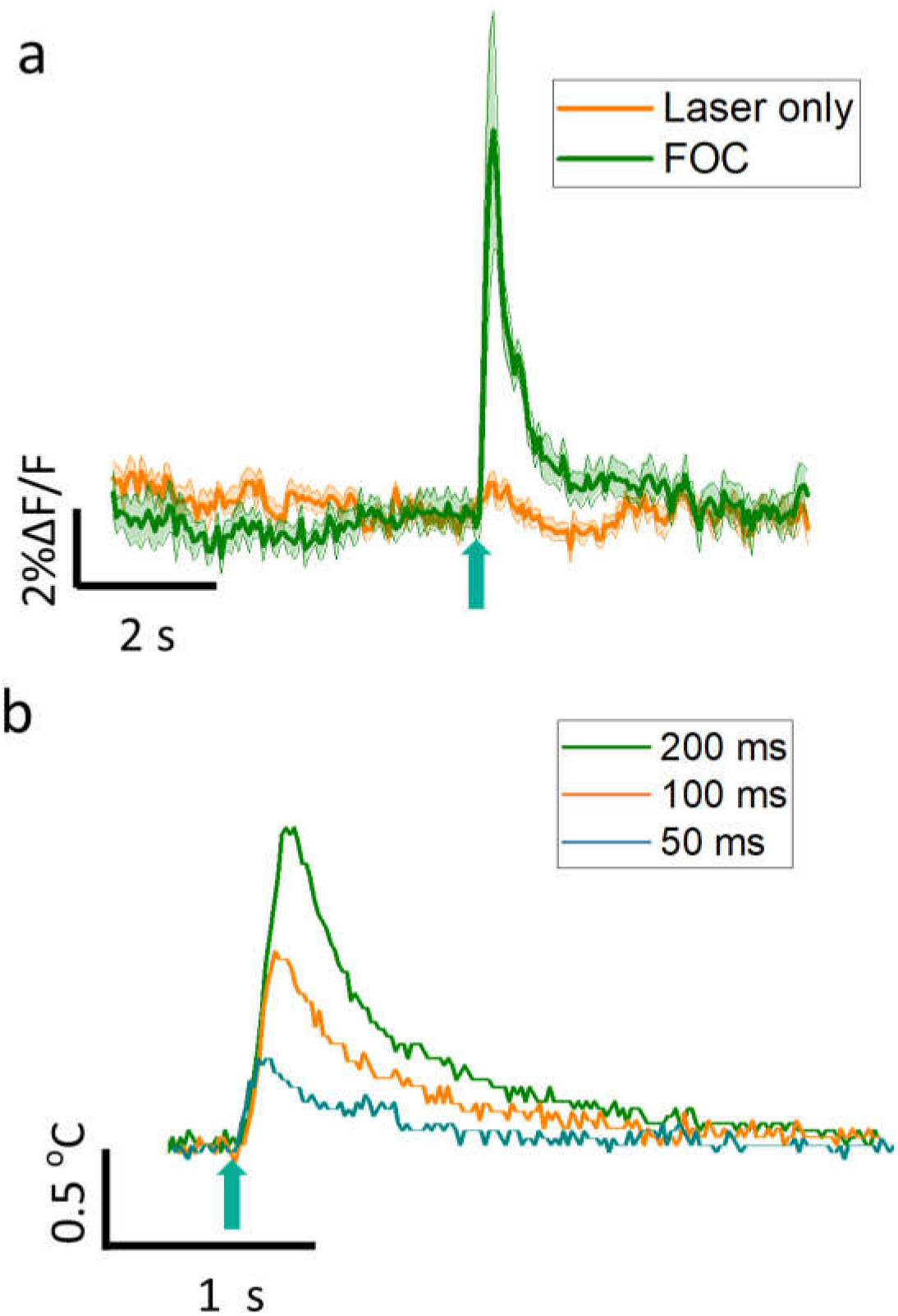
FOC-mediated neural activation is not induced by laser or heat. **a**, Average trace of neuronal calcium trace in response to laser stimulation (n=13) and FOC stimulation (n=12). Shaded area: ± SD. **b**, Surface temperature dynamics of FOC tip during laser excitation.

### FOC induces direct activation of the targeted cortical area in mouse brain without the involvement of the auditory pathway

Since we demonstrated that the FOC can reliably induce neural activation with high spatial precision in vitro, we next moved on to investigate whether FOC can successfully induce neural activation in vivo in mouse brain, with similar spatial precision. The mouse was deeply anesthetized, and a cranial window was made above the primary somatosensory cortex (S1) and primary auditory cortex (A1) (**Fig. 5a**). First, we investigated whether FOC can produce activation of the local cortical area. The FOC was brought close to contact the brain surface. Laser pulses with 200 ms and 50 ms duration were delivered to the FOC, and neural activities were recorded with a tungsten electrode. We observed robust local field potential (LFP) response to the FOC stimulation for both stimulation durations, with response latency of 15.87 ± 1.34 ms (n=3) (**Fig. 5b**), which is indicative of direct neural activation. When we lifted the FOC up by 100 μm without contacting the brain or immersing ACSF, the FOC failed to induce any neural activation (**Fig. 5c**). This result indicates that the neural activation is induced by optoacoustic waves that have minimal propagation in the air. Next, we delivered FOC stimulation to the ipsilateral A1 region, which is approximately 2 mm away from the S1 recording site and failed to detect any neural responses (**Fig. 5c**), demonstrating superior spatial confinement of the FOC stimulation in vivo. Since auditory stimulation by ultrasound has been reported^12,13^, we examined whether the auditory pathway is involved in the FOC stimulation. One cranial window was made above the S1 region, and another on the contralateral A1region. The FOC stimulation was delivered to the S1 region, and the recording electrode was placed in the ipsilateral S1 or contralateral A1 region (**Fig. 5d**). If the auditory pathway is involved, we would observe strong responses in the contralateral A1 with ∼50 ms delay ^13^. However, a FOC stimulation of 200 ms duration on the S1 evoked robust LFP response on ipsilateral S1, but failed to evoke any response in the contralateral A1 (**Fig. 5e**). Histological examination showed no damage of the neural tissue at the optoacoustic stimulation site. Finally, to rule out the possibility of laser or ultrasound-induced electrical artifact of the electrode, we record voltage change on the FOC surface in saline and found that laser pulses of 200 ms duration produced no voltage change on the FOC tip (**Fig. 5f**). We repeated the same measurements on three different mice and obtained the same results. Collectively, these data suggest that the FOC produces direct neural stimulation in vivo with high spatial and temporal precision, without the involvement of the auditory pathway.

**Figure 5.**
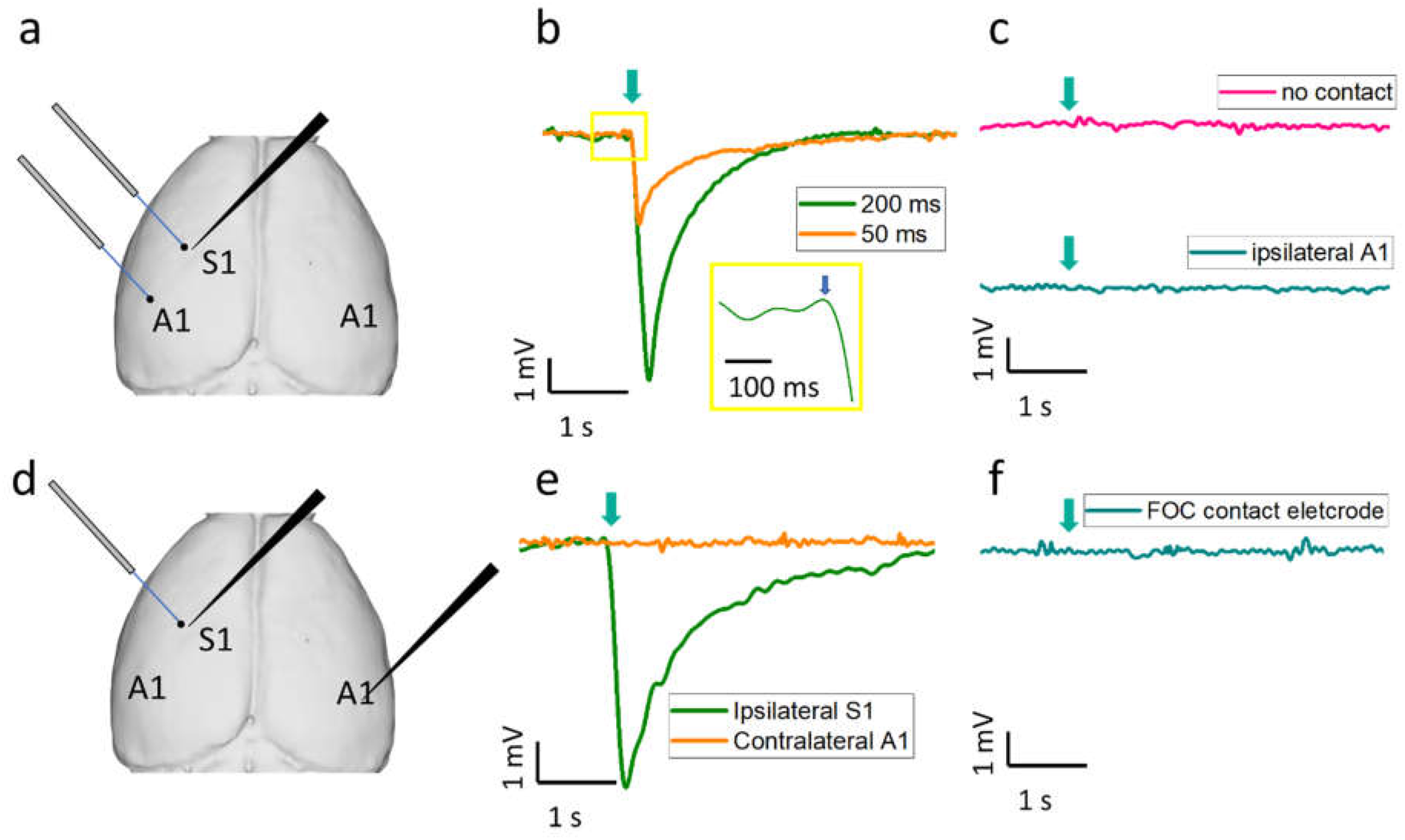
FOC induces direct and localized neural activation in vivo in mouse brain. **a**, Placement of FOC in S1 and A1, and ipsilateral recording electrode in S1 test the spatial confinement of FOC stimulation. **b**, S1 LFP response to 50 ms and 200 ms FOC stimulation delivered to S1. Insert: zoomed in LFP trace showing response latency. **c**, S1 LFP response to FOC stimulation delivered to A1 and S1 LFP response to FOC stimulation delivered to S1 without contact. **d**, Placement of FOC in S1 and recording electrode in ipsilateral S1 and contralateral A1 to test the involvement of the auditory pathway. **e**, LFP response pf ipsilateral S1 and contralateral A1 to S1 FOC stimulation. **f**, Voltage response to the FOC stimulation when electrode contacts the FOC. Green arrow: stimulation onset.

## Discussion

We demonstrated a miniaturized FOC that can induce neural activation with high spatial precision both in vitro and in vivo. The FOC has a ball-shaped coated tip with a diameter of ∼600 μm and allows omnidirectional generation of optoacoustic wave at 1 to 5 MHz. Fast attenuation of acoustic intensity is achieved by the nano-composite diffusion layer and the ball-shaped geometry. The neural response is shown to be neither thermal nor laser-induced.

An important observation is that our FOC system directly activates targeted cortical area in vivo, instead of indirect activation through the auditory pathway. This finding is supported by multiple pieces of evidence. First, the FOC is able to stimulate cortical neurons in culture, where no auditory circuits are involved; Second, the FOC stimulated targeted cortical area only with less than 20 ms delay, which is indicative of direct stimulation; Third, the stimulation was delivered to the cortex directly, avoiding any possible bone transduction to the cochlear; Finally, the FOC stimulation failed to induce neural response on the auditory cortex contralateral to the stimulation site, thus eliminating the involvement of the auditory pathway. Our results are consistent with previous studies where focused ultrasound can directly stimulate hippocampal slices, isolated retina in vitro, and activate mouse M1 in vivo with less than 50 ms delay^5,14,15^. Yet, Guo et al. and Sato et al. argued that transcranial ultrasound neural stimulation could reach the cochlear pathway first, which leads to downstream activation of the auditory cortex and additional cortical areas^12,13^. One of the reasons for this controversy is the wide range of ultrasound parameters used in these studies and additional studies are needed to further characterize the contribution of these two pathways under different ultrasound stimulation parameters.

Although the FOC and piezo-based transducer both can generate acoustic waves in the ultrasonic frequency, significant differences exist between these two devices. First, the FOC with a diameter of around 600 μm is significantly smaller than most commercially available ultrasound transducers. The FOC size can be further reduced by using optical fiber with a smaller diameter and reducing the coating layers. The much smaller size allows the FOC to be implantable and can be used for behavior study in live free-running animals, which is impossible with traditional ultrasound transducers. Second, the FOC generate 2-microsecond pulsed acoustic waves repeated at 3.6 KHz. Thus, the duty cycle of our optoacoustic wave is about 0.72%. This low duty cycle avoids ultrasound heating of biological tissues, making the FOC device particularly suitable for in vivo applications. Third, for most transcranial ultrasound neural modulation applications, the peak pressure ranges from 0.1-2 MPa^1,2^, and recently Guo et al. demonstrated neural modulation with acoustic pressure as low as 250 kPa^13^. The peak acoustic pressure generated by the FOC was measured to be ∼0.1 MPa which falls into the range of ultrasound intensity used for neural modulation reported in the literature. Fourth, a wide range of frequencies has been reported to achieve neural modulation (200 kHz to 32 MHz)^1,2^. Generally, the lower frequency is used for transcranial stimulation, and a higher frequency is used to achieve high spatial confinement. The current FOC generates broadband ultrasound wave with multiple peaks ranging from 0.5 to 5 MHz, where the spatial precision is achieved because the optoacoustic wave from the FOC tip attenuates quickly.

The mechanism of optoacoustic neural stimulation is yet to be investigated ^1^. Ultrasound neural stimulation and optoacoustic neural stimulation are similar in the way that they both create a mechanical disturbance on the neuronal membrane and are likely to share the same mechanism. Two mechanisms are proposed for ultrasound neural stimulation: ultrasound induced intramembrane cavitation ^30-32^ and activation of mechanosensitive channels ^33,34^. Future studies using specific mechanosensitive channels blockers are needed to identify the relative contributions of these two mechanisms.

Finally, we note that the FOC tip is versatile and can be easily customized for more advanced applications. The propagation of the generated optoacoustic wave depends largely on the FOC tip geometry. The ball-shaped FOC tip allows omnidirectional optoacoustic wave propagation and fast attenuation, while other geometries can be adopted to generate forward, focused and even patterned, complex acoustic field^35-37^, which can be used for neural modulation at even higher spatial resolution. Additionally, the fiber-based design allows the FOC to be implanted for longitudinal behavior study in live animals. Given the increasing popularity of ultrasound neuromodulation, the compactness, cost-effectiveness, and versatility of FOC open a lot of opportunities to utilize the optoacoustic effect to achieve high-precision neural stimulation. Without the need for genetic modification, we expect that FOC will eventually be used for neural modulation on human subjects, similar to electrode-based deep brain stimulation but in a metal-free manner.

## Methods

### Fabrication of FOC

The FOC was fabricated by first coating the fiber with a light diffusion layer, followed by coating of an absorption layer. ZnO nanoparticles (Sigma-Aldrich) were mixed with epoxy at a concentration of 15% by weight, and a polished multimodal optic fiber with 200 μm core diameter (Thorlabs) was dipped into the mixture and quickly pulled out. After 30 min curing at room temperature, the diffusion layer was coated on the fiber tip. The absorption layer was fabricated by dipping of the diffusion layer coated fiber into a graphite powder and epoxy mixture (30% by weight), quickly pulled put and cure in room temperature. This process was repeated to ensure the absorption layer has enough thickness to absorb all the photons.

### Acoustic wavefront mapping

For mapping of the acoustic wavefront, an EKSPLA OPO Laser with pulse width 5 ns, petition rate 10 Hz was coupled into the FOC fiber as excitation laser. Photoacoustic signals were acquired in a water tank by a low-frequency transducer array (L7-4, PHILIPS/ATL) and processed by an ultrasound imaging system (Vantage128, Verasonics Inc.).

### Primary neuronal and glial cultures

Primary cortical neuron cultures were derived from Sprague-Dawley rats. Briefly, cortices were dissected out from embryonic day 18 (E18) rats of either sex and then digested with papain (0.5 mg/mL in Earle’s balanced salt solution) (Thermofisher scientific) and plated on poly-D-lysine coated coverslips. For primary neuron cultures, cells were first plated in Dulbecco's Modified Eagle Medium (Thermofisher scientific) containing 10% fetal bovine serum (Thermofisher scientific) and 1% GlutaMAX^tm^ (Thermofisher scientific), which was then replaced 24 hours later by a feeding medium (Neurobasal medium supplemented with 2% B-27 (Thermofisher scientific) and 1% GlutaMAX^tm^ (Thermofisher Scientific). Thereafter, the medium was replaced every 3 to 4 days until use. For primary glial cultures, cells were cultured in Dulbecco's Modified Eagle Medium (Thermofisher scientific) containing 10% fetal bovine serum (Thermofisher scientific) and medium was replaced every 3 to 4 days.

### Calcium imaging

Oregon Green™ 488 BAPTA-1 AM (Invitrogen) was dissolved in 20% Pluronic F-127 in DMSO at a concentration of 1 mM as stock solution. Before imaging, cells were incubated with 2 μM OGB-1 for 30 min, followed by incubation with normal medium for 30 min. During imaging, cells were placed in extracellular solution for cortical neurons containing 150 mM NaCl, 4 mM KCl, 10 mM HEPES, 10 mM glucose, 2 mM CaCl_2_ (pH 7.4). Calcium imaging was performed on a lab-built inverted fluorescence microscope, with a LED at 470 nm as excitation light source, an emission filter (FBH520-40, Thorlabs), an excitation filter (MF469-35, Thorlabs) and a dichroic mirror (DMLP505R, Thorlabs). Image sequences were acquired with a scientific CMOS camera (Zyla 5.5, Andor) at 20 frames per second.

### Animal surgery

Adult (age 14-16 weeks) C57BL/6J mice were used. Mice were initially anesthetized using 5% isoflurane in oxygen and then placed on a standard stereotaxic frame, maintained with 1.5 to 2 % isoflurane. Toe pinch was used to determine the level of anesthesia throughout the experiments and body temperature was maintained with a heating pad. The hair and skin on the dorsal surface targeted brain regions were trimmed. Craniotomies were made on primary somatosensory (S1) (AP −1.34 ML 2.25) and primary auditory cortex (A1) (AP −2.46 ML 4.25) based on stereotaxic coordinates using a dental drill and artificial cortical spinal fluid was administrated to immerse the brain. After stimulation and recordings, the mice were perfused with saline and 10% Formalin, and the brain was removed and sectioned for histology.

### Local field potential recording

Electrophysiology was performed using tungsten microelectrodes (0.5 to 1 MΩ; Microprobes). Tungsten microelectrodes were driven to recording sites through cranial windows (r = 1.5 mm) based on stereotactic coordinates and confirmed by electrophysiological signatures. The electrodes were positioned with a micromanipulator (Siskiyou). Extracellular recordings were acquired using a Multi Clamp 700B amplifier (Molecular Devices), filtered at 0.1 to 100 Hz, and digitized with an Axon DigiData 1550 digitizer (Molecular Devices). For calculation of response latency, the pre-stimulation period in each recording was used to obtain baseline mean and SD. The threshold was determined by mean ± 2*SD. The latency was determined when the voltage crosses the threshold for the first time.

### Data analysis

Calcium images were analyzed using ImageJ. The fluorescence intensity was measured by selecting the soma. Optoacoustic waveforms, calcium traces, temperature traces, and electrophysiological traces were analyzed using Origin. Data shown are mean ± SD.

## Acknowledgments

The authors thank Y. Bai for help with building the fluorescence microscope, S. Cha with the testing on in vivo experiments. We also thank the Boston University Experimental Pathology Laboratory Service Core for help with histology experiments. This work was supported by R01 NS109794 to J.X.C. to X.H.

## Author contributions

JXC conceived the concept of using optoacoustic waves for neurostimulation; Y.J. and L.L. designed and fabricated the FOC device; Y.J., H.J.L. and L.L. performed the experiments; Y.J., H.J.L., H.T., C.Y., X.H., and J.C. discussed and analyzed the data; H.M. provided neuron cultures; Y.J. and J.C. wrote and all authors discussed and edited the manuscript; X.H. and J.C. supervised the project.

